# Perturbing RNA localization for functional study in neurons

**DOI:** 10.1101/2025.04.05.646707

**Authors:** Mengting Han, Maylin L. Fu, Yanyu Zhu, Alexander A. Choi, Emmy Li, Jon Bezney, Sa Cai, Leanne Miles, Yitong Ma, Stanley Qi

## Abstract

Spatial RNA organization plays a pivotal role in diverse cellular processes and diseases, but the functional implications of spatial RNA localization remain underexplored. We present CRISPR-mediated transcriptome <u>o</u>rganization (CRISPR-TO) that harnesses RNA-guided, nuclease-dead dCas13 for programmable control of RNA localization in live cells. CRISPR-TO enables targeted localization of RNAs to diverse subcellular compartments, including p-bodies, stress granules, telomeres, and nuclear stress bodies, across cell types. In primary cortical neurons, we demonstrate that repositioned mRNAs undergo local translation along neurites and at neurite tips and co-transport with ribosomes, with *β*-actin mRNA localization enhancing the formation of dynamic filopodial protrusions and inhibiting axonal regeneration. Furthermore, CRISPR-TO-enabled parallel screening in primary neurons identifies *Stmn2* mRNA localization as a driver of neurite outgrowth. By enabling large-scale perturbation of the spatial transcriptome, CRISPR-TO bridges a critical gap left by current sequencing and imaging technologies, offering a versatile platform for high-throughput functional interrogation of RNA localization in living cells and organisms.

---

Spatial RNA localization is a fundamental mechanism that orchestrates the spatiotemporal organization and regulation of macromolecules, playing a critical role in RNA metabolism and protein synthesis (1). Since the initial discovery of the asymmetric distribution of actin mRNA in ascidian embryos (2), distinct patterns of RNA organization have been identified across various genes and organisms. Examples include the localization of β-actin mRNA to the growth cone of retinal cells in Xenopus (3) and the polarized distribution of various mRNAs in the intestinal epithelium of mice (4). The biological significance of RNA organization is particularly pronounced in large, polarized cells such as neurons. By positioning specific RNAs at the right time and place, cells effectively bridge the vast scale gap between nanometer-sized molecules and cells that are six to nine orders of magnitude larger (5). Moreover, aberrant RNA localization has been correlated with an increasing number of neurological diseases, including amyotrophic lateral sclerosis (ALS), fragile X syndrome (FXS), and spinal muscular atrophy (SMA) (1, 6).

High-throughput imaging (*e*.*g*., MERFISH and seq-FISH) and sequencing methods (*e*.*g*., APEX-seq and ExSeq) have uncovered the spatial localization of thousands of RNAs within distinct subcellular compartments across diverse life forms (7-11). Despite these advances, the functional roles of the spatial transcriptome remain largely unexplored, with functional insights limited to a small subset of mRNAs (1). This gap arises due to major challenges in perturbing endogenous RNA localization in primary cells like neurons.

Traditional approaches for perturbing RNA organization involve complex genome engineering, such as removing specific elements that control RNA localization (12). However, the localization elements for most RNAs remain unidentified, and such deletions can inadvertently affect RNA stability, splicing, and translation (13). Approaches that tag RNAs with MS2 repeats bound by the MS2 coat protein (MCP) fused to a subcellular compartment protein have allowed for the study of specific RNAs (14, 15). However, these approaches are labor intensive, challenging for large-scale studies, and less feasible in primary cells. To address these challenges, we develop a novel, programmable method for perturbing endogenous RNA localization, termed CRISPR-mediated transcriptome organization (CRISPR-TO), enabling functional investigation of endogenous RNA localization in diverse living cells.

Class 2 type VI CRISPR-Cas13 systems have been identified as RNA-guided ribonucleases in mammalian cells (16-18). Catalytically dead dCas13 variants exhibit efficient and specific RNA binding without cleavage (17-21). Leveraging these programmable features, we developed CRISPR-TO by coupling dCas13 with subcellular compartment-specific signals or motor proteins via a chemical-inducible ABI-PYL1 dimerization system (22).

For cytoplasmic RNA localization, we fused PYL1 with structural proteins of of p-bodies (23), or stress granules (24). CRISPR-TO showed efficient localization of *GAPDH* mRNA to these compartments (**Figure 1A-B**). For nuclear RNA localization, we fused PYL1 with a core component of the telomere nucleoprotein complex (25), or a marker protein of nuclear stress body (26). We targeted telomeric repeat-containing RNA (*TERRA*), a long noncoding RNA (lncRNA) (27). We observed efficient *TERRA* lncRNA at telomeres after ABA treatment (**Figure 1C**). We also confirmed efficient localization of reporter mRNA to nuclear stress bodies (**Figure 1D**). In addition to passive diffusion-mediated localization, we extended CRISPR-TO for active RNA transport along microtubules. By attaching PYL1 to truncated variants of kinesin for anterograde or retrograde transport (28), we directed endogenous *GAPDH* mRNA localization towards the plus or minus ends of microtubules (**Figure 1E-F**). In summary, our findings highlight the versatility of CRISPR-TO in localizing both mRNA and noncoding RNA to diverse subcellular compartments via either passive diffusion or active transport mediated by motor proteins.

**Figure 1.**
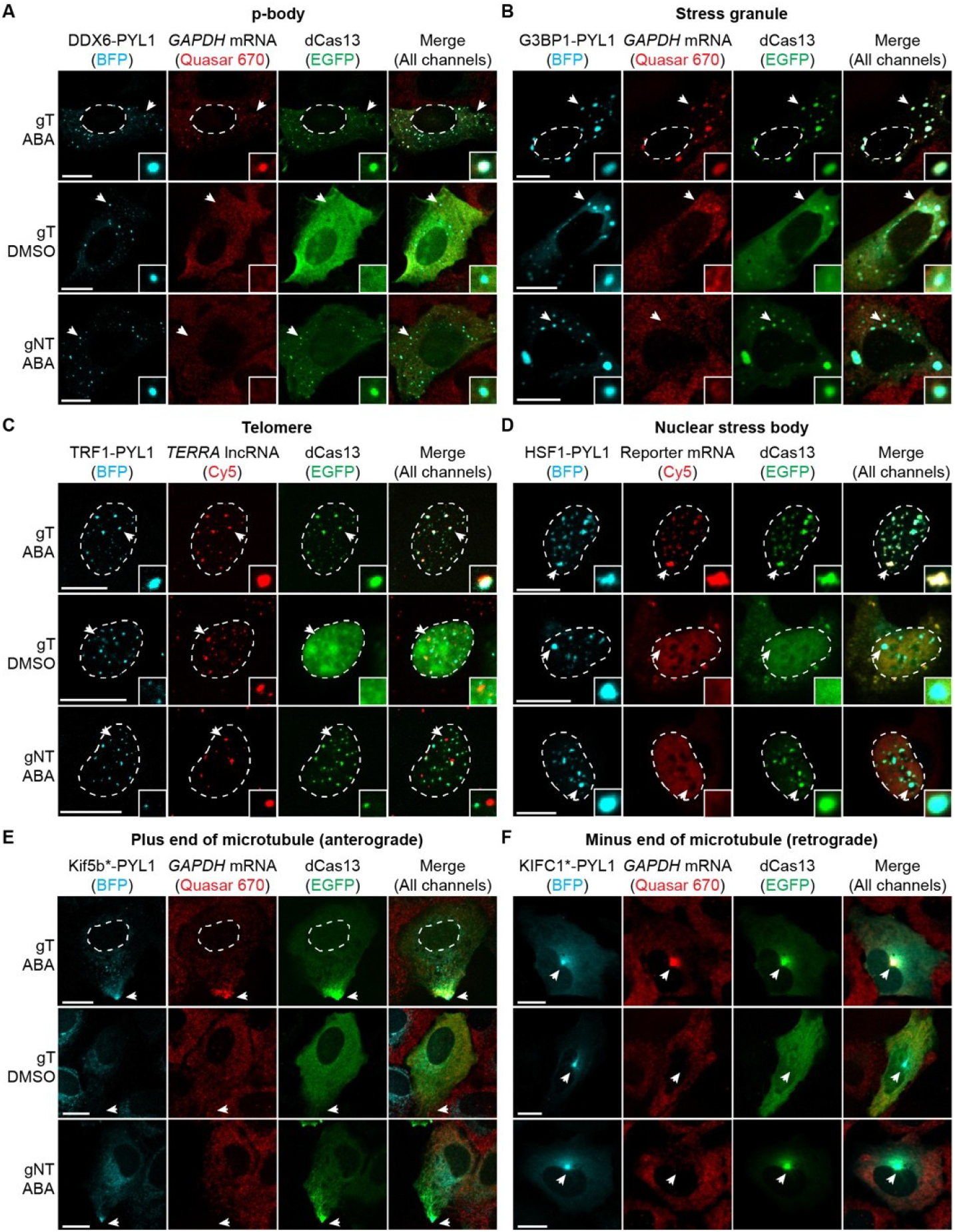
CRISPR-TO can efficiently localize mRNA and ncRNA to different subcellular compartments. **A-B**: Representative images showing CRISPR-TO-mediated localization of endogenous *GAPDH* mRNA to p-bodies (**A**) and stress granules under stress conditions (**B**) under different gRNA and ABA conditions. Inset images represent magnification of the region indicated by the white arrow. **C-D**: Representative images showing CRISPR-TO-mediated localization of *TERRA* lncRNA to telomeres (**C**) and GCN4 reporter mRNA to nuclear stress bodies under stress conditions (**D**) under different gRNA and ABA conditions. **E-F**: Representative images showing CRISPR-TO-mediated localization of *GAPDH* mRNA at the leading edge of the cell (**E**) or centrosome (**F**) under different gRNA and ABA conditions. The shape of the nucleus is depicted with a white dotted line in cells that are hard to visualize the nucleus. Scale bar, 20 µm.

We utilized CRISPR-TO to manipulate the spatial localization of endogenous RNAs in primary neurons. We transfected mouse primary cortical neurons with plasmids encoding CRISPR-TO components and a membrane-localized fluorescent protein (CAAX-Crimson) to visualize neurite morphology (29). After CRISPR-TO perturbation for 24 hours, dCas13 protein and *Actb* mRNA were transported to neurite tips, both near (200 µm, inset 1) or far (716 µm, inset 3) from the soma (**Figure 2A-B**). We also captured numerous mRNA particles along the neurites (inset 2). Quantification of the fluorescence intensity revealed a 10-fold increase in endogenous *Actb* mRNA localization at neurite tips after ABA treatment compared to DMSO (**Figure 2C**).

**Figure 2.**
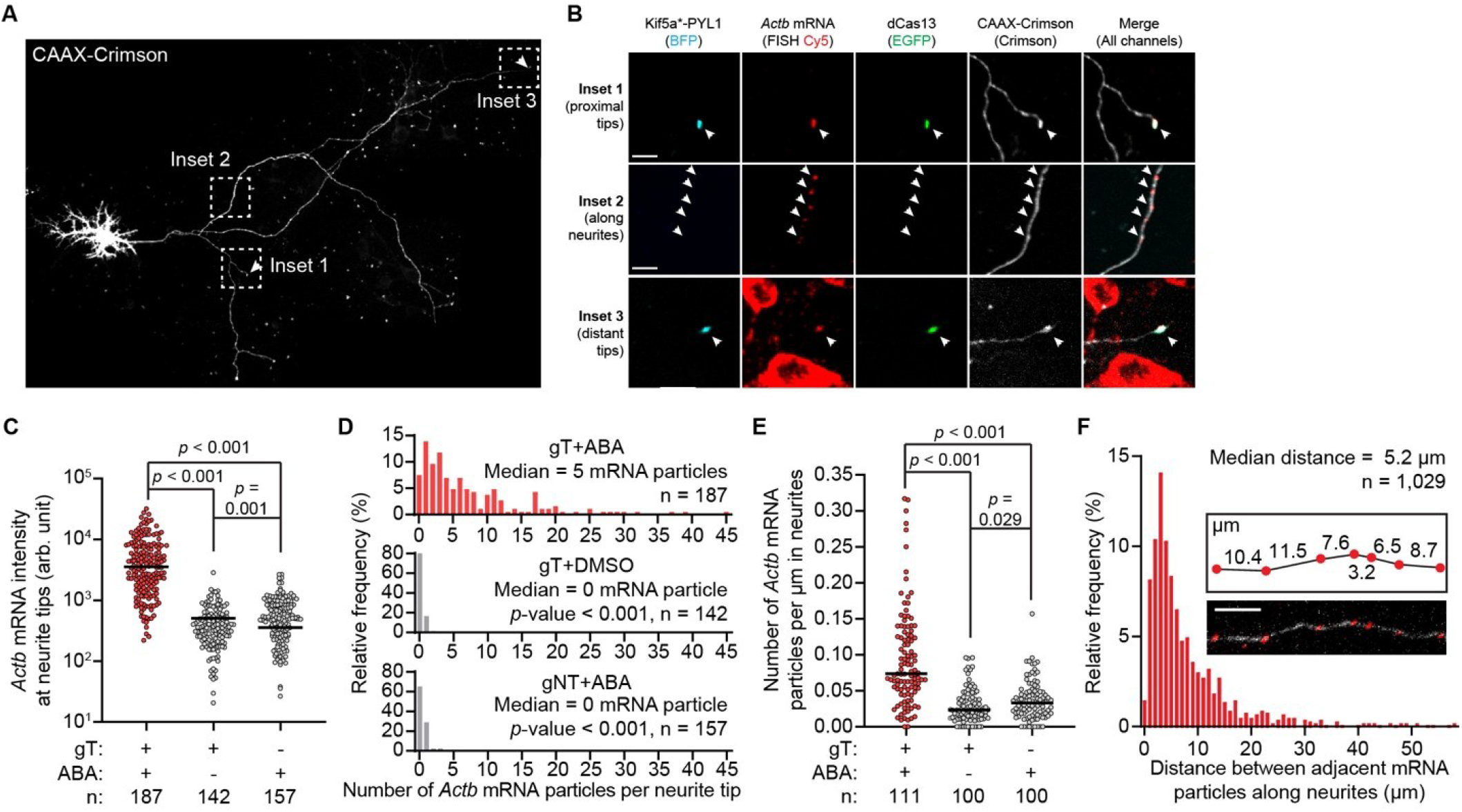
Efficient control of endogenous mRNA transport over ultralong distances in primary neurons via CRISPR-TO. **A-B:** Representative images of a neuron (**A**, scale bar, 100 µm) and the insets of its neurite and neurite tips (**B**, scale bar, 10 µm) in the gT+ABA group. gT, gRNAs targeting *Actb* mRNA. **C**: Quantification of the fluorescence intensity of endogenous *Actb* mRNA in the neurite tips. n indicates the number of quantified neurite tips for each group. **D:** Frequency distribution of the number of endogenous *Actb* mRNA particles in one neurite tip for the gT+ABA, gT+DMSO, and gNT+ABA groups. *p*-value was calculated using unpaired t-test between gT+ABA group and indicated groups. **E**: Quantification of the number of endogenous *Actb* mRNA particles per µm in neurites. n indicates the number of quantified neurite segments for each group. **F**: Distribution of the distance between two endogenous *Actb* mRNA particles in the neurites for gT+ABA. The inset figures show an example image (the bottom panel) with *Actb* mRNA particles (red) distributed along a fragment of a neurite (white) in the gT+ABA group and the distance between two adjacent *Actb* mRNA particles (the top panel). Scale bar, 10 µm.

Further analysis showed that each neurite tip contained a median of five *Actb* mRNA particles following perturbation, compared to 0 particle per neurite tip for control groups (**Figure 2D**). The number of *Actb* mRNA particles varied, with some tips containing up to 45 particles. We quantified the density of *Actb* mRNA particles along neurites and observed a median of 0.07 particle per µm (**Figure 2E**), with a median distance of 5.2 µm between adjacent particles (**Figure 2F**). Compared to the DMSO group, ABA treatment increased the number of transporting *Actb* mRNA particles along neurites by 3.2-fold (**Figure 2E**).

We next investigated whether mRNA localized to neurite tips by CRISPR-TO can be locally translated. To test this, we transfected primary mouse cortical neurons with plasmids encoding CRISPR-TO, CAAX-HaloTag, and the reporter mRNA, followed by ABA or DMSO treatment for 6 hours. After UV conversion, we monitored Dendra2 green fluorescence recovery at neurite tips for 50 minutes. In the ABA group, green fluorescence increased steadily, while it decreased in the DMSO group (**Figure 3A-B**). The mean fluorescence recovery over the period across 72 neurite tips in the ABA group was significantly higher than the DMSO group (**Figure 3C**), indicating local translation.

**Figure 3.**
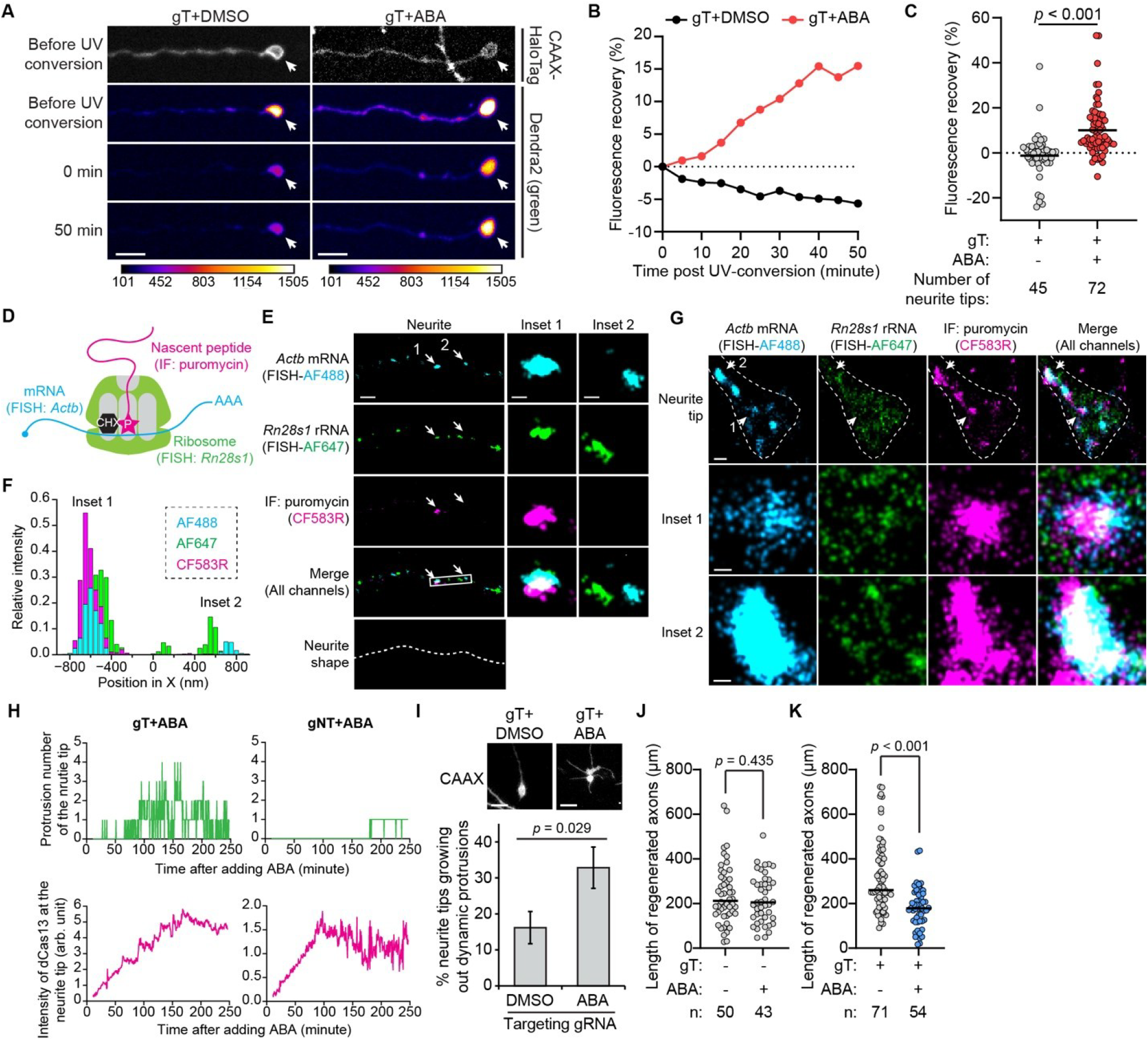
Functional characterization of mRNA localization on local translation, neurite morphology, and regeneration. **A-B**: Representative confocal images (**A**) and quantification (**B**) of green fluorescence recovery of Dendra2 in the neurite tip of gT+ABA group is higher than that of gT+DMSO group. Scale bar, 10 µm. Calibration bar indicates the green fluorescence intensity of Dendra2. **C**: Quantification of the green fluorescence recovery of Dendra2 at 50 minutes post UV-conversion. Data was presented as means ± SEM. Unpaired t-test. **D**: Schematics showing the labeling of *Actb* mRNA via RNA FISH, ribosome via RNA FISH staining of *Rn28s1* rRNA, and translation site via ribopuromycylation (31). CHX: cycloheximide. P: puromycin. **E**: Representative super-resolution images showing the three-color staining of *Actb* mRNA, ribosome, and translation site inside neurites for the gT+ABA group after CRISPR-TO perturbation. Scale bar, 1 µm. Inset images represent magnification of the region pointed by the white arrow. Scale bar, 200 nm. **F**: Histogram of the relative intensity of the white box region in **E. G**: Representative super-resolution images showing the three-color staining of *Actb* mRNA, ribosome, and translation site inside neurite tips for the gT+ABA group after CRISPR-TO perturbation. Scale bar, 500 nm. Inset images represent magnification of the region pointed by the white arrow. Scale bar, 100 nm. The shape of neurite tips is depicted with a white dotted line. **H**: The number of protrusions and the intensity of dCas13 at the neurite tip of gT+ABA and gNT+ABA group as a function of ABA treatment time. **I**: Representative microscopic images showing the morphology of two types of neurite tips (top panel; scale bar, 10 µm) and percentage of neurite tips growing out dynamic protrusions (bottom panel) after DMSO or ABA treatment in neurons with gT (n = 66-68 neurite tips per group). Error bars represent SEM calculated according to Bernoulli distributions. Two-sided Fisher’s exact test. **J-K**: Quantification of the lengths of regenerated axons under different conditions. Unpaired t-test. n indicates the number of quantified regenerated axons for each group.

To confirm ribosome co-transport and local translation of endogenous mRNA along neurites and at neurite tips, we performed three-color stochastic optical reconstruction microscopy (STORM) super-resolution imaging (30) to visualize *Actb* mRNA, ribosomes, and active translation sites (31) in the gT+ABA group (**Figure 3D**). Our super-resolution images revealed *Actb* mRNA particles colocalized with both ribosome and puromycin signals in neurites, indicating active translation during transport (**Figure 3E-F**, inset 1). Notably, fewer puromycin-labeled sites compared to *Actb* mRNA particles and ribosomes suggest that only a subset of transported mRNAs is actively translated. Similarly, aggregated *Actb* mRNA clusters colocalized with ribosomes and active translation sites at neurite tips (**Figure 3G**), indicating local translation.

The dendritic and axonal localization of β-actin mRNA is known to regulate axon branch formation, growth cone guidance, and dendritic filopodia density, primarily studied by blocking its localization with antisense oligonucleotides or overexpressing neurite-localized exogenous β-actin mRNA (3, 32, 33). However, the direct functional influence of endogenous β-actin mRNA in neurites has not been thoroughly analyzed. Given that β-actin polymerization drives sensory filopodia protrusion (34), we hypothesized that recruiting endogenous β-actin mRNA to neurite tips might increase filopodial protrusions.

To test this, we applied CRISPR-TO in primary mouse cortical neurons to transport β-actin mRNA to neurite tips and monitored morphology changes following ABA addition. Initially, neurite tips exhibited a spindly and stable structure. After 90 minutes of ABA treatment, multiple (1 to 4) dynamic protrusions emerged. In contrast, the gNT control group showed no new protrusions despite dCas13 enrichment at neurite tips in both groups (**Figure 3H**). We monitored ∼70 neurites tips over 7 hours and observed 33% developed dynamic protrusions after ABA treatment, significantly higher than the 16% observed with DMSO (**Figure 3I**).

We next examined whether β-actin mRNA localization influences axon regeneration using a microfluidic device-based axonal injury model that separates axons from the soma (35). With gNT, no difference in axon regeneration was observed with or without ABA, suggesting that ABA itself does not affect axonal growth (**Figure 3J**). In the presence of gT, we observed significantly shorter axonal lengths after ABA treatment compared to untreated neurons (**Figure 3K**), suggesting that promoting β-actin mRNA localization inhibits axon regeneration. These findings reveal a relationship between β-actin mRNA localization and changes in neurite morphology.

The programmability of CRISPR-TO enables high-throughput genetic screens for the functional study of spatial RNA localization on a large scale. As a proof-of-concept, we performed an arrayed, paralleled screen using CRISPR-TO to track the functional influence of neurite localization of 21 mRNAs on neurite outgrowth in primary mouse cortical neurons (**Figure 4A**). Considering the possible role of local mRNA translation at neurite tips, we expected that a subset of these mRNAs would have measurable effects on neurite outgrowth upon perturbation via CRISPR-TO.

**Figure 4.**
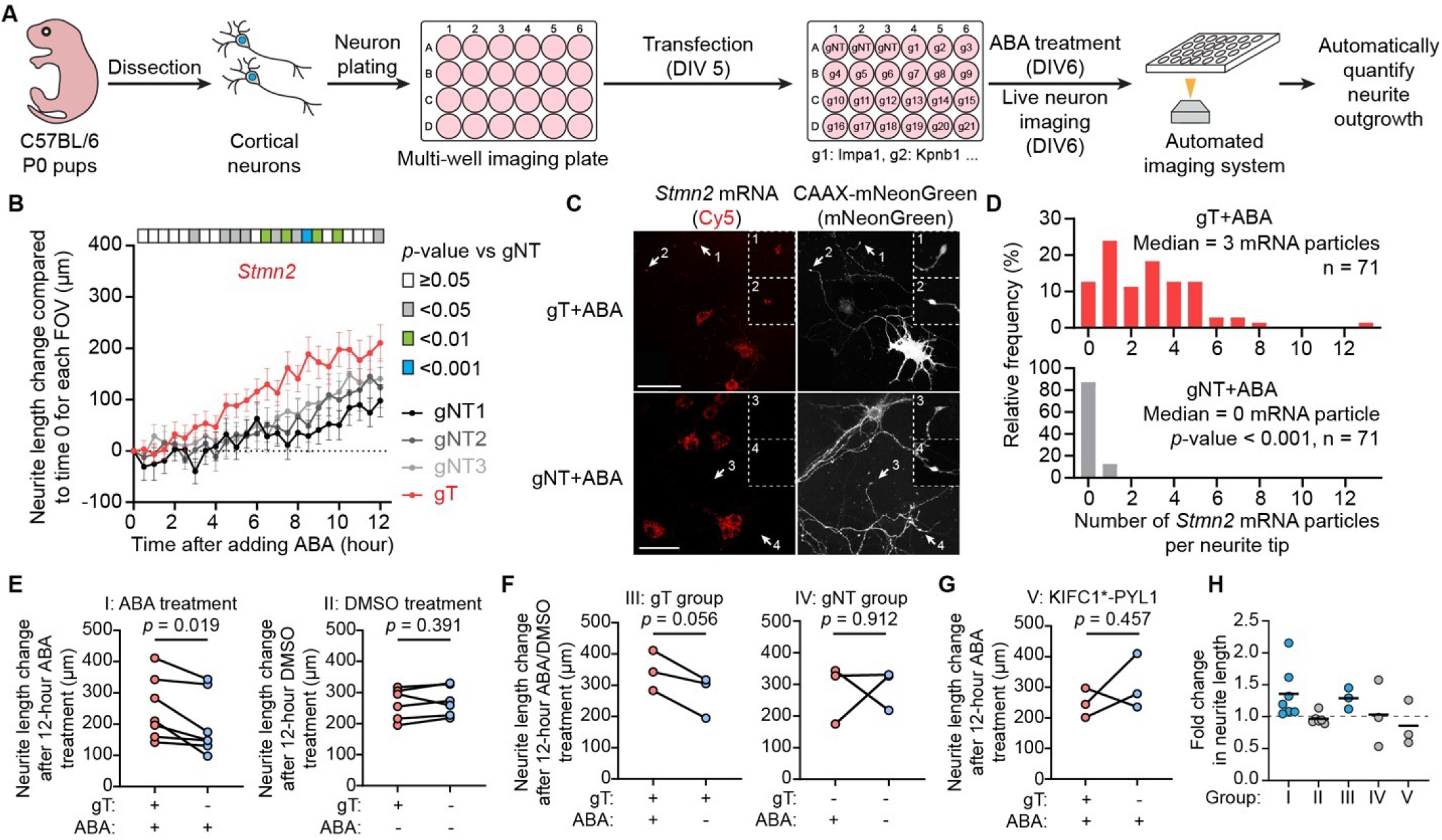
CRISPR-TO screens in primary neurons reveal mRNAs whose neurite localization promotes neurite outgrowth. **A**: Schematics of applying CRISPR-TO to screen for mRNAs whose neurite localization influences neurite outgrowth using high-content imaging system. **B**: Quantification of the neurite length changes over time after adding ABA. Mean ± SEM of different fields of views is shown for each time point. The color bar above each plot shows the *p*-value of each time point compared with the gNT group. **C**: Representative confocal images showing endogenous *Stmn2* mRNA enriched at neurite tips after CRISPR-TO perturbation. Scale bar, 50 µm. Inset images surrounded by the dotted line represent magnification of the region indicated by the white arrow. **D**: Frequency distribution of the number of endogenous *Stmn2* mRNA particles in one neurite tip for the gT+ABA and gNT+ABA groups. p-value was calculated using unpaired t-test between the two groups. **E-F**: Neurite length change after 12-hour ABA or DMSO treatment for neurons expressing CRISPR-TO components, CAAX-mScarlet-I, and gRNAs targeting *Stmn2* or gNT. **G**: Neurite length change after 12-hour ABA treatment for neurons expressing CRISPR-TO components, CAAX-mScarlet-I, and gRNAs targeting *Stmn2* or gNT. The *p*-value was calculated using paired t-test. **H**: Fold change of the left group compared to the right group for each pair in **E-G**. The black bar indicates the mean value.

We cultured neurons in multi-well plates and transfected plasmids expressing CRISPR-TO components, a neurite tracing marker (CAAX-mScarlet-I), and gRNAs targeting candidate mRNAs or non-targeting controls. After transfection, we treated neurons with ABA or DMSO and imaged every 30 minutes for 12 hours using an automated high-content microscope to track neurite outgrowth.

We compared the neurite outgrowth of each of 21 mRNA candidates to the three gNT groups. Among the 21 mRNAs screened, 4 mRNAs showed significant effects on neurite outgrowth (p < 0.01) at specific time points (**Figure 4B**), and were selected as hits.

One hit mRNA, *Stmn2*, encodes Stathmin-2, a tubulin-binding protein involved in microtubule dynamics (36), axonal regeneration(37, 38), and ALS pathology as a promising therapeutic target (39). *Stmn2* mRNA has been identified in axons and growth cones via microarrays (40, 41), but the functional relevance of its mRNA localization was unclear. It is possible that localized *Stmn2* mRNA at neurite tips can exhibit local translation to facilitate stabilizing microtubules and promoting axonal regeneration (42). To validate *Stmn2* mRNA localization as a screen hit, we performed RNA FISH. Without CRISPR-TO perturbation (gNT+ABA), *Stmn2* mRNA was found in both cell body and soma-proximal neurites (**Figure 4C**). CRISPR-TO perturbation (gT+ABA) resulted in *Stmn2* mRNAs enrichment at neurite tips, with a median of 3 mRNA particles per tip compared with zero in the gNT+ABA group (**Figure 4D**).

We repeated the neurite outgrowth assay under various conditions to validate the functional role of *Stmn2* mRNA localization. After 12 hours of ABA treatment, neurons in the gT group showed significantly greater neurite length increases compared to the gNT group, whereas DMSO treatment had no significant effect (**Figure 4E**). In addition, ABA treatment specifically increased neurite length in the gT group, but not in the gNT group (**Figure 4F**). In contrast, the retrograde transport did not significantly affect neurite outgrowth (**Figure 4G**), possibly because most *Stmn2* mRNAs are naturally located in the soma of *in vitro* cultured primary mouse cortical neurons. In conclusion, our findings demonstrate the neurite localization of *Stmn2* mRNA promotes neurite outgrowth (**Figure 4H**).

CRISPR-TO offers several advantages over traditional methods for perturbing the spatial transcriptome. It bypasses the need for genomic alterations, as dCas13 directly binds endogenous RNA via a complementary gRNA. It enables RNA localization to various subcellular compartments by modifying the localization signal fused with PYL1. CRISPR-TO is effective in primary cells like neurons, where spatial transcriptomes are crucial and closely linked to pathological processes (1, 6). Its programmability allows for high-throughput functional screening of the spatial transcriptome by varying gRNA sequences to target different RNAs. CRISPR-TO-mediated perturbation of spatial transcriptome should provide valuable insights into the biological functions of spatial RNA organization across diverse cell types and disease progresses.

